# False facts and false views: coalescent analysis of truncated data

**DOI:** 10.1101/006445

**Authors:** Einar Árnason

## Abstract

Darwin’s dictum on false facts and false views points the way to opening the road to truth via cogent criticism of the published record. Here I discuss a case in which a truncated dataset (false facts) is used for coalescent analysis of historical demography that reaches a foregone conclusion of a bottleneck of numbers (false views).

---

“False facts are highly injurious to the progress of science, for they often endure long; but false views, if supported by some evidence, do little harm, for everyone takes a salutary pleasure in proving their falseness; and when this is done, one path towards error is closed and the road to truth is often at the same time opened” (Darwin, 1871, p. 385). Darwin’s dictum is in full force and I apply it here to a case where false facts have led to false views hoping to open the road to truth.

Ólafsdóttir *et al*. (2014) studied demographic history of Atlantic cod, *Gadus morhua*, at Iceland using mtDNA isolated from vertebrae from archaeological sites. They compare their results to already published results from modern times (citing Árnason, 2004). They notice a reduction in haplotype and nucleotide diversity in modern times and use coalescent analysis to infer a bottleneck of numbers at 1400–1500 and a marked reduction of effective population size, *N*_*e*_, in modern times. They use Approximate Bayesian Computation, ABC, to model three population size scenarios evaluated by matches to summary statistics.

A key problem of the study of Ólafsdóttir *et al*. (2014) is the handling of the data of the modern samples for which they cite Árnason (2004) which summarizes data from several papers on variation of cytochrome *b* from various localities in the Atlantic ocean. The primary data on Iceland are not in that paper. Árnason *et al*. (2000) published the original primary data on Icelandic cod, a paper not cited by Ólafsdóttir *et al*. (2014).

First, the numbers reported in their Table S3 and said to represent “Modern frequency” are not in accordance with the original correct data (Árnason et al., 2000; Árnason, 2004) (Table 1). The original data have 519 individuals with 23 segregating sites defining 30 haplotypes (Table III of Árnason *et al*., 2000) whereas Table S3 reports different numbers for common and rare haplotypes and total numbers and omits many haplotypes. There are discrepancies for many but not all haplotypes (Table 1). There also are discrepancies between the numbers for modern times reported in Table 2 of the paper and in supplemental Table S3: sample size of 503 vs 499, number of haplotypes 10 vs 8, with 7 vs 6 segregating sites.

**Table 1.**
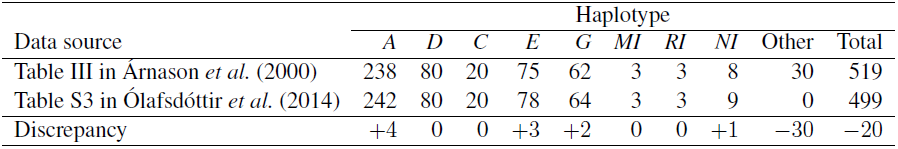
Discrepancies in frequencies of haplotypes in data for modern-times. First row is from Table III in Árnason et al. (2000) Árnason et al. (2000). Second row is truncated data from Table S3 of Ólafsdóttir *et al*. (2014) Ólafsdóttir *et al*. (2014) said to be modern-times data from Árnason (2004) Árnason (2004). Third row is discrepancy added (+) and ommitted (−) between the first two rows. Other represents a pool of 22 rare haplotypes omitted in Ólafsdóttir *et al*. (2014).

| Data source | Haplotype |  |  |  |  |  |  |  |  | Total |
| --- | --- | --- | --- | --- | --- | --- | --- | --- | --- | --- |
|  | *A* | *D* | *C* | *E* | *G* | *MI* | *RI* | *NI* | Other |  |
| Table III in Árnason *et al.* (2000) | 238 | 80 | 20 | 75 | 62 | 3 | 3 | 8 | 30 | 519 |
| Table S3 in Ólafsdóttir *et al.* (2014) | 242 | 80 | 20 | 78 | 64 | 3 | 3 | 9 | 0 | 499 |
| Discrepancy | +4 | 0 | 0 | +3 | +2 | 0 | 0 | +1 | −30 | −20 |

Second, Ólafsdóttir *et al*. (2014) do not use all the data of the modern sample (Árnason *et al*., 2000). They truncate the data by omitting 22 haplotypes, all singleton (17), doubleton (3), one triplet and one quadruplet haplotype. These truncations of the original data result in a dataset of 499 individuals with 8 haplo-types and 6 segregating sites (Table S3). They are false facts. Coalescent analysis in general proceeds by tracing the ancestry of a sample to a common ancestor.

By its nature coalescence is sensitive to the size and composition of a sample. If a real sample from a natural population in true fact was both large (as the modern sample Árnason *et al*., 2000) and had few or no rare alleles (as in Table S3 Ólafsdóttir *et al*., 2014) the genealogy would be characterized by long internal and few or no external branches. There would be a deficiency of low frequency variants and an excess of middle frequency variants. This would be a clear sign of a declining population under coalescence theory (Wakeley, 2009, page 120). Using the truncated data dataset for the Bayesian skyride plot (Minin *et al*., 2008) under BEAST (Drummond *et al*., 2012) stacks the odds and Ólafsdóttir *et al*. (2014) reach a foregone conclusion of a population bottleneck and low effective size in the modern times. These are false views.

The 1500–1550 and the 1910 samples stand out from the rest (Table S3 Ólafsdóttir *et al*., 2014) and also influence the skyride analysis. The 1500–1550 sample has a relatively large number of haplotypes and segregating sites, a relative evenness in haplotype frequencies giving high nucleotide diversity (*π̂* = 0.0059 compared to *π̂* = 0.0052 the modern sample Árnason *et al*. (2000), and *π̂* = 0.0047 for the truncated data in Table S3 Ólafsdóttir *et al*. (2014)). The 1910 sample has few haplotypes and segregating sites, a relatively high frequency of the most common haplotype and consequently low nucleotide diversity (*π̂* = 0.0043). Nucleotide diversity estimates the scaled effective population size *θ* = 2*N*_*e*_*µ* Wakeley (2009). These divergent samples along with the truncated dataset of the modern sample are drivers of the apparent bottlenecks in skyride analysis Ólafsdóttir *et al*. (2014).

I have generated distributions of the number of segregating sites, the number of haplotypes and the nucleotide diversity from 1000 random samples of size 36 representing the sample size of the 1500–1550 sample and of 1000 samples of size 23 representing the 1910 sample of Ólafsdóttir *et al*. (2014) by random sampling from the Árnason *et al*. (2000) dataset. At least 25% of the distributions had a greater number of segregating sites than 6 and a greater number of haplotypes than 8 reported for the 1500–1550 sample in Table S3 (Ólafsdóttir *et al*., 2014). More than 7% had a higher nucleotide diversity than the 1500–1550 sample. For the 1910 sample 3 out of 1000 had equal or fewer segregating sites than the sample, about 6% had fewer or equal numbers of haplotypes and 25% had a lower nucleotide diversity. Thus these divergent samples are within sampling errors of the modern haplotype frequencies (Árnason *et al*., 2000). Therefore, the bottle-necks (Ólafsdóttir *et al*., 2014) are spurious resulting from a combination of the use of the truncated modern-times data and sampling variation in the small ancient samples.

There also are internal discrepancies between results given in Table 2 and in supplemental Table S3 of Ólafsdóttir *et al*. own data. For example, Table 2 reports 9 haplotypes and 7 segregating sites for the 1500–1550 sample. However, the detailed data reported in Table S3 are 8 haplotypes defined by 6 segregating sites (number of segregating sites can be determined from Table III of Árnason *et al*. (2000) or from Figure 1 of Árnason (2004)). Similarly, there should be 5 and not 4 segregating sites in the 1650–1700 dataset and 3 and not 4 segregating sites in the 1910 dataset of Ólafsdóttir *et al*. (2014).

Third, ABC analysis in general proceeds by simulating random datasets and selecting a small subset of these that are most similar to the real dataset based on congruence of summary statistics. Ólafsdóttir *et al*. (2014) used number of haplotypes and number of segregating sites and summary statistics based on these in their analysis. Discrepancies in summary statistics described above may bias the selection of the sub-samples of 500 out of a million random datasets. Also they report type I and II errors of 44% and 46% for a scenario of two bottlenecks compared to a scenario of a single bottleneck or a constant population size. The statement that “the ABC analysis supported the scenario of two bottlenecks over the scenario of either a single bottleneck or constant population size…” is strange given the very high type I and type II errors rates.

Fourth, the method section of the paper seems to imply that all the molecular work was done in a dedicated ancient DNA laboratory in Canada. However, the supplement states that only DNA isolation was done in dedicated ancient DNA laboratory in Canada and that the rest of the molecular work from PCR amplification to sequencing was done in a lab in Reykjavik where “no previous work on Atlantic cod had taken place”. However, this statement is inaccurate. The post PCR work was actually done in shared facilities where Atlantic cod DNA of modern samples, both mitochondrial and nuclear *Pan* I (Árnason, 2004; Árnason *et al*., 2009; Eiríksson & Árnason, 2013), has been amplified and sequenced for many years. It is, therefore, not clear how established criteria for ancient DNA work (Cooper & Poinar, 2000) were adhered to.

Also, there is no mention of how it was determined that the vertebrae sampled from archaeological sites represent vertebrae from different individuals. How, for example, can we know that the high evenness and high nucleotide diversity of the 1500–1550 sample or the low diversity of the 1910 sample is not pseudo-replication due to sampling multiple vertebrae from the same individual?

